# Identification of a novel HIF-1α-α_M_β_2_ Integrin-NETosis axis in fibrotic interstitial lung disease

**DOI:** 10.1101/2020.01.03.894196

**Authors:** Akif A. Khawaja, Deborah L.W. Chong, Jagdeep Sahota, Charis Pericleous, Vera M. Ripoll, Helen L. Booth, Saif Khan, Manuel Rodriguez-Justo, Ian P. Giles, Joanna C. Porter

## Abstract

Neutrophilic inflammation correlates with mortality in fibrotic interstitial lung disease (ILD) however, the underlying mechanisms remain unclear. We aimed to determine whether aberrant neutrophil activation is a feature of ILD and the relative role of hypoxia. We used lung biopsies and bronchoalveolar lavage (BAL) from ILD patients to investigate the extent of hypoxia and neutrophil activation in lungs of patients with ILD. We complemented these findings with *ex vivo* functional studies of neutrophils from healthy volunteers to determine the effects of hypoxia. We demonstrate for the first time HIF-1α staining in neutrophils and endothelial cells in ILD lung biopsies. Hypoxia enhanced both spontaneous and phorbol 12-myristate 13-acetate (PMA)-induced neutrophil extracellular trap (NET) release (NETosis), neutrophil adhesion, and trans-endothelial migration. Hypoxia also increased neutrophil expression of the α_M_ and α_X_ integrin subunits. Interestingly, NETosis was induced by α_M_β_2_ integrin activation and prevented by cation chelation. Finally, NETs were demonstrated in the BAL from ILD patients, and quantification showed significantly increased cell-free DNA content and MPO-citrullinated histone H3 complexes in ILD patients compared to non-ILD controls. Our work indicates that HIF-1α upregulation may augment neutrophil recruitment and activation within the lung interstitium through activation of β_2_ integrins. Our results identify a novel HIF-1α-Integrin-NETosis axis for future exploration in therapeutic approaches to fibrotic ILD.

## Introduction

The interstitial lung diseases (ILD) are a group of diffuse parenchymal lung disorders that can result in pulmonary fibrosis (PF)^1^. Despite recent advances in diagnostics and therapeutics, ILD is still associated with substantial morbidity and mortality^2^. Neutrophil activation may play a role in ILD and in particular in the most severe fibrotic form, idiopathic PF (IPF). The pathogenesis of IPF is unknown but is thought to involve an aberrant response to repetitive epithelial injury, with associations to genes and proteins linked to epithelial function, integrity and repair. Progressive epithelial damage, and aberrant wound repair leads to extensive scar formation and correlates, clinically, with worsening hypoxia. Increasing desaturation during exercise^3^ or sleep^4^ is a significant predictor of mortality. Further evidence from animal models suggests that hypoxia may actually contribute to a vicious cycle of disease progression^5^. This has led to the view that hypoxia itself may contribute to worsening of PF but how this occurs is not known.

Hypoxia, a state in which oxygen supply is inadequate for tissue demands, modulates gene expression via transcriptions factors called hypoxia inducible factors (HIF). There are 3 members of the HIF family, HIF-1α, HIF-2α and HIF-3α, which bind conserved DNA sequences or Hypoxia Response Elements (HRE). Although it seems plausible that the IPF lung is hypoxic much of the evidence is indirect. Levels of lactic acid, a metabolite generated in response to hypoxia, are high in IPF lung tissue supporting the concept of a hypoxic microenvironment^6^ and HIF-1α and 2α have been shown, *ex vivo*, to be expressed in lung biopsies from patients with IPF, in some but not all reports^7,8^. Additional genomic studies in IPF patients show up-regulation of hypoxia-related gene signatures, including TGF-β^9^, the key fibrotic cytokine in PF, and of the HIF-1α pathways^8,10^.

The contribution of neutrophils to ILD has also been relatively less studied compared to other inflammatory and fibrotic diseases. Early studies began to explore the potential role of neutrophils in IPF^11–14^, however research focus has since shifted to other cell types. The number of neutrophils in the bronchoalveolar lavage (BAL) has been shown to predict both disease severity in IPF^15^ and the development of PF in patients with hypersensitivity pneumonitis^16^. In addition, neutrophil extracellular traps (NETs) have been shown to indirectly drive PF by stimulation of collagen production from fibroblasts *in vitro*^17^, and NET release (NETosis) has been associated with PF in older mice *in vivo*^18^ with loss of peptidyl arginine deiminase (PAD)-4, a key neutrophil enzyme for NETosis, being protective^19^. Neutrophils are also associated with disease severity in acute lung injury and acute respiratory distress syndrome (ARDS)^20,21^ however, their precise contribution remains uncertain^22^. Neutrophil depletion can ameliorate disease features in mouse models of ARDS^23^ and a reduction in neutrophil infiltration^24^, or knock-down of neutrophil elastase (NE) attenuates fibrosis in bleomycin-induced mouse models of PF^25^. Taken together, these studies implicate a contributory role of neutrophils to fibrotic ILD.

Neutrophil survival is a tightly regulated process. Prolonged survival can delay resolution of inflammation and can cause damage to surrounding cells and tissues; however, if apoptosis is premature, antimicrobial function can be compromised^26^. Hypoxia drives neutrophil survival via HIF-1α-dependent NF-κB activation^27^. In addition, HIF-2α has also been shown to be important in regulating neutrophil function^28^. Few reports address the effects of hypoxia upon NETosis. Inhibition of HIF-1α can reduce NET release^29^, whilst pharmacological stabilisation of HIF-1α increases phagocyte bactericidal activity^30^, implicating a role for down-stream targets of HIF-1α in leukocyte function.

Given the importance of hypoxia and HIF signalling in neutrophil function and the emerging role of neutrophils as key drivers of pulmonary disease, we sought evidence for hypoxia and NETosis in the lungs of patients with ILD and the functional effects of low oxygen levels upon *ex vivo* neutrophil function and activation.

## Results

### Patient demographics

BAL was obtained from 11 ILD patients and 7 non-ILD controls undergoing diagnostic bronchoscopies. Demographics, clinical history and treatments at the time of sample collection are listed in Table 1. Within the ILD cohort: 4 (36%) had IPF, 3 (27%) had nonspecific interstitial pneumonia, 3 (27%) had chronic hypersensitivity pneumonitis (HP) and 1 (10%) had unclassifiable ILD. Our non-ILD control group underwent diagnostic bronchoscopy due to: 5 (71%) investigation of haemoptysis, 1 (14.5%) right middle lobe collapse and 1 (14.5%) previous tracheal schwannoma patients undergoing yearly bronchial surveillance. Only the ILD group had lung function tests, as part of standard patient care. None of the patients recruited were taking anti-fibrotic drugs at the time of bronchoscopy. The differential cell population obtained from BAL from patients are listed in supplementary table 1.

**Table 1:** Patient clinical and demographic data. Clinical details were recorded for all subjects at the time of bronchoalveolar lavage. Our ILD cohort had a mean age of 69±5.9 years and consisted of 4 patients with IPF, 3 patients with fibrotic NSIP, 3 patients with chronic HP and 1 patient with unclassifiable ILD. Our non-ILD control group had a mean age of 51±10.1 years and consisted of 5 patients with haemoptysis, 1 patient with RML collapse and 1 patient with tracheal schwannoma. Lung function was only obtained for the ILD cohort as part of standard care. Abbreviations: FVC, forced vital capacity; HP, hypersensitivity pneumonitis; ILD, interstitial lung disease; IPF, idiopathic pulmonary fibrosis; NSIP, nonspecific interstitial pneumonia; RML, right middle lobe; TLCO, transfer factor for carbon monoxide.

|  |  | ILD | Non-ILD Control |
| --- | --- | --- | --- |
| Cohort size (n) |  | 11 | 7 |
| Age (years ±SD) | | 69 ±5.9 | 51 ±10.1 |
| Sex (M:F) |  | 8:3 | 4:3 |
| Smoking Status (current:ever:never) |  | 2:6:3 | 1:2:4 |
| Clinical Features |  |  |  |
| Diagnosis |  | IPF (4), fibrotic NSIP (3), chronic HP (3), unclassifiable ILD (1) | Haemoptysis (5), RML collapse (1), tracheal schwannoma (1) |
| Forced Vital Capacity (FVC) (litres) (mean ±SD) [% predicted] | | 2.7 ±0.9 [80.4 ±17.6%] | - |
| O<sub>2</sub> saturation (mean ±SD) | | 96.2 ±1.7% | - |
| TLCO (mmol/min/kPa) (mean ±SD) | | 50.8 ±12.3 | - |
| Current Medications |  |  |  |
| Corticosteroids | Inhaled | Budesonide/formoterol fumarate dihydrate (1), fluticasone propionate/salmeterol xinafoate (1) | Budesonide/formoterol fumarate dihydrate (1) |
|  | Oral | Methyl-prednisolone (1), prednisolone (1) | Prednisolone (1) |
| Nonsteroidal anti-inflammatory drugs |  | Aspirin (4) | Naproxen (1) |

### Neutrophils and endothelial cells stain positive for HIF-1α in the ILD lung

Given reports of localised hypoxia in pulmonary disease^31^, we examined ILD lung biopsy samples for evidence of hypoxia. Initial HIF-1α staining demonstrated positive staining in the endothelium and polymorphonuclear cells, with very little staining in the fibrotic interstitium and overlying epithelium and no staining in control sections (fig. 1A, B). As NETosis has been implicated in several immunopathologies, we also stained lung sections for myeloperoxidase (MPO) and NE (fig. 1C, D), highlighting the presence of neutrophils within the pulmonary vasculature. Taken together, this staining pattern suggests that tissue-specific hypoxia and neutrophil recruitment may be a feature of the ILD lung. These findings led us to examine the effects of hypoxia upon neutrophil function.

**Figure 1:**
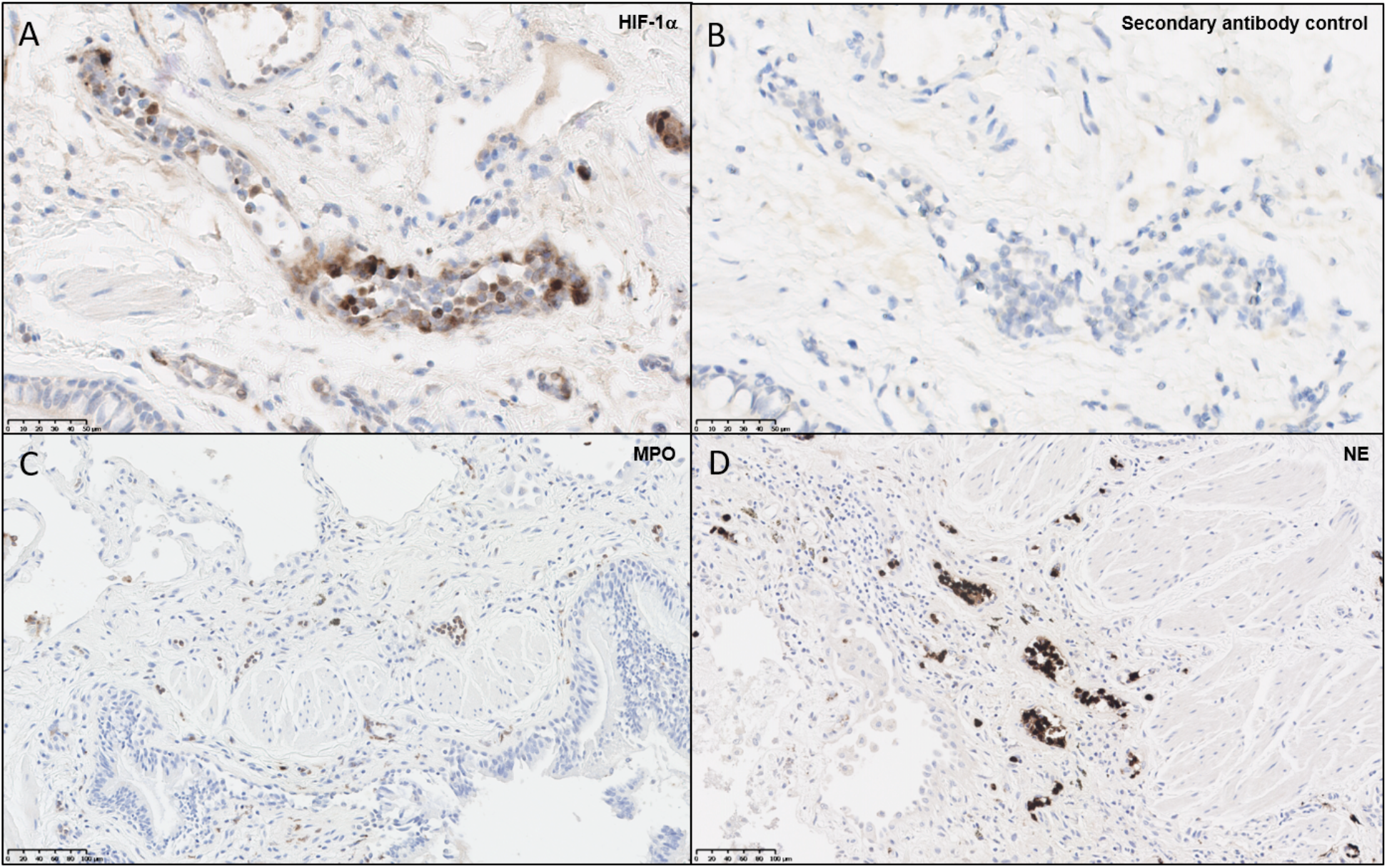
Neutrophils and endothelial cells express HIF-1α within the ILD lung. Paraffin-embedded ILD lung biopsies were cut and stained for immunohistochemical evidence of hypoxia and neutrophil infiltration. **(A)** Slides stained for HIF-1α displayed positive brown staining within microvascular endothelial cells and polymorphonuclear cells, whilst **(B)** secondary antibody controls did not display positive staining. To verify whether neutrophils were present in the ILD lung, additional stains were performed for **(C)** MPO and **(D)** NE, both of which displayed positive brown stains within blood vessels. Abbreviations: HIF-1α, hypoxia-inducible factor 1α; MPO, myeloperoxidase; NE, neutrophil elastase.

### Hypoxic exposure does not affect hydrogen peroxide generation but promotes NETosis

Pharmacological HIF-1α stabilisation has been reported to enhance bacterial killing and NETosis^29,30^, however, these studies were performed using atmospheric oxygen levels. We therefore assessed for any alteration in function, described below, of healthy neutrophils under normoxia (21% oxygen) and hypoxia (1% oxygen). First, we verified hypoxia by examining neutrophil cell lysates for the presence of HIF-1α and HIF-2α at various time points. We observed rapid stabilisation of HIF-1α under hypoxia, with delayed HIF-2αstabilisation (fig. 2A).

**Figure 2:**
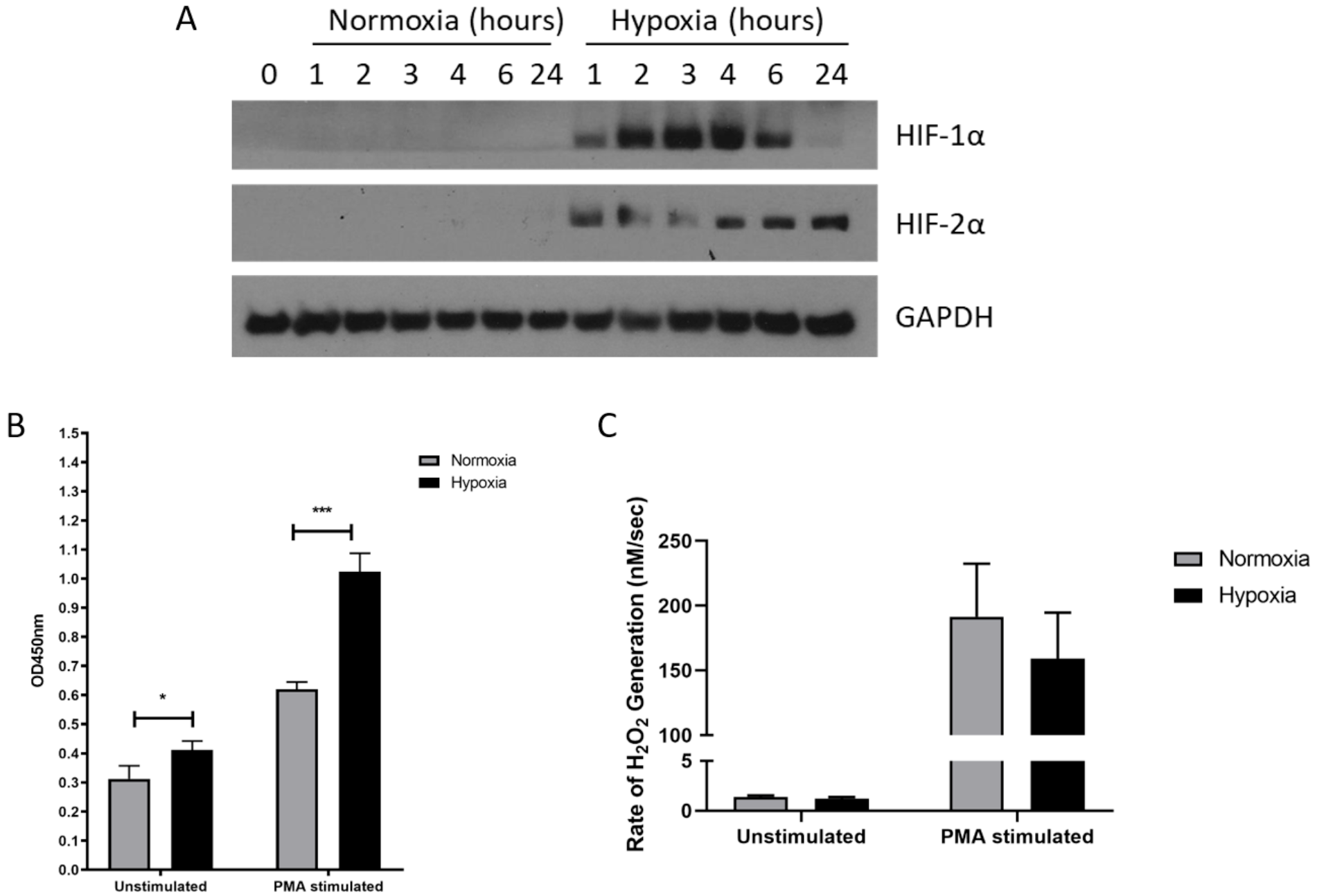
Hypoxia enhances NETosis but not hydrogen peroxide production. The effects of hypoxia upon neutrophil activation was first assessed. **(A)** The induction of hypoxia was verified by Western blot, probing for HIF-1α and HIF-2α. **(B)** NETosis was then evaluated by capture ELISA, which detects MPO-citrullinated histone H3 complexes. Data are presented as the mean and SEM from three independent experiments and analysed by two-way ANOVA with a Dunnett’s multiple comparison test. **(C)** Hydrogen peroxide generation was examined using Amplex^®^ UltraRed in absence and presence of 50nM PMA. Data are presented as the mean and SEM neutrophils isolated from 7 different donors and analysed by two-way ANOVA with a Dunnett’s multiple comparison test. *= p<0.05, ***= p<0.001. Abbreviations: HIF, hypoxia-inducible factor; PMA, phorbol 12-myristate 13-acetate.

Having demonstrated induction of hypoxia, we then assessed neutrophil supernatants for MPO-citrullinated histone H3 complexes, which are specific for NETs. Hypoxic neutrophils displayed greater levels of both spontaneous (+1.31-fold, p<0.05) and phorbol 12-myristate 13-acetate (PMA)-induced (+1.65-fold, p<0.001) NETosis (fig. 2B). As reactive oxygen species generation is thought to drive NETosis^32,33^, we also examined hydrogen peroxidase (H_2_O_2_) production. In these experiments however, rates of H_2_O_2_ generation were comparable between oxygen states for both unstimulated (1.4±0.1nM/sec vs. 1.2±0.2nM/sec) and PMA-stimulated (191.7±40.83nM/sec vs. 159.3±35.51nM/sec) neutrophils (fig. 2C).

### Neutrophil adhesion and trans-endothelial migration are enhanced under hypoxia

Having found an effect on NETosis, we next examined integrin activation and neutrophil adhesion, which are also implicated in NET release^34,35^. We measured neutrophil adhesion to primary human endothelial cells in the absence or presence of PMA (a general integrin activator) or lipopolysaccharide (LPS) (to mimic infectious stimuli), stimuli that activate NETosis via distinct pathways^36^. Hypoxia increased both unstimulated (23.6±4.0% vs 2.7±1.6%, p<0.05) and LPS-stimulated (35.7±4.8% vs 11.3±1.4%, p<0.05) adhesion to resting endothelium, whilst PMA-stimulated adhesion, which was already high, was unaffected (fig. 3A-C). We then looked at adhesion to endothelium pretreated with TNF-α, to mimic an inflammatory event. Whilst unstimulated neutrophil adhesion to TNF-α activated endothelial cells was not altered by hypoxia, there was a 3.22- and 2.11-fold increase in PMA- (21.2±6.3%vs 68.1±8.4%. p<0.05) and LPS-stimulated (23.2±2.8%vs 49.0±2.3%, p<0.05) adhesion respectively (fig. 3D-F).

**Figure 3:**
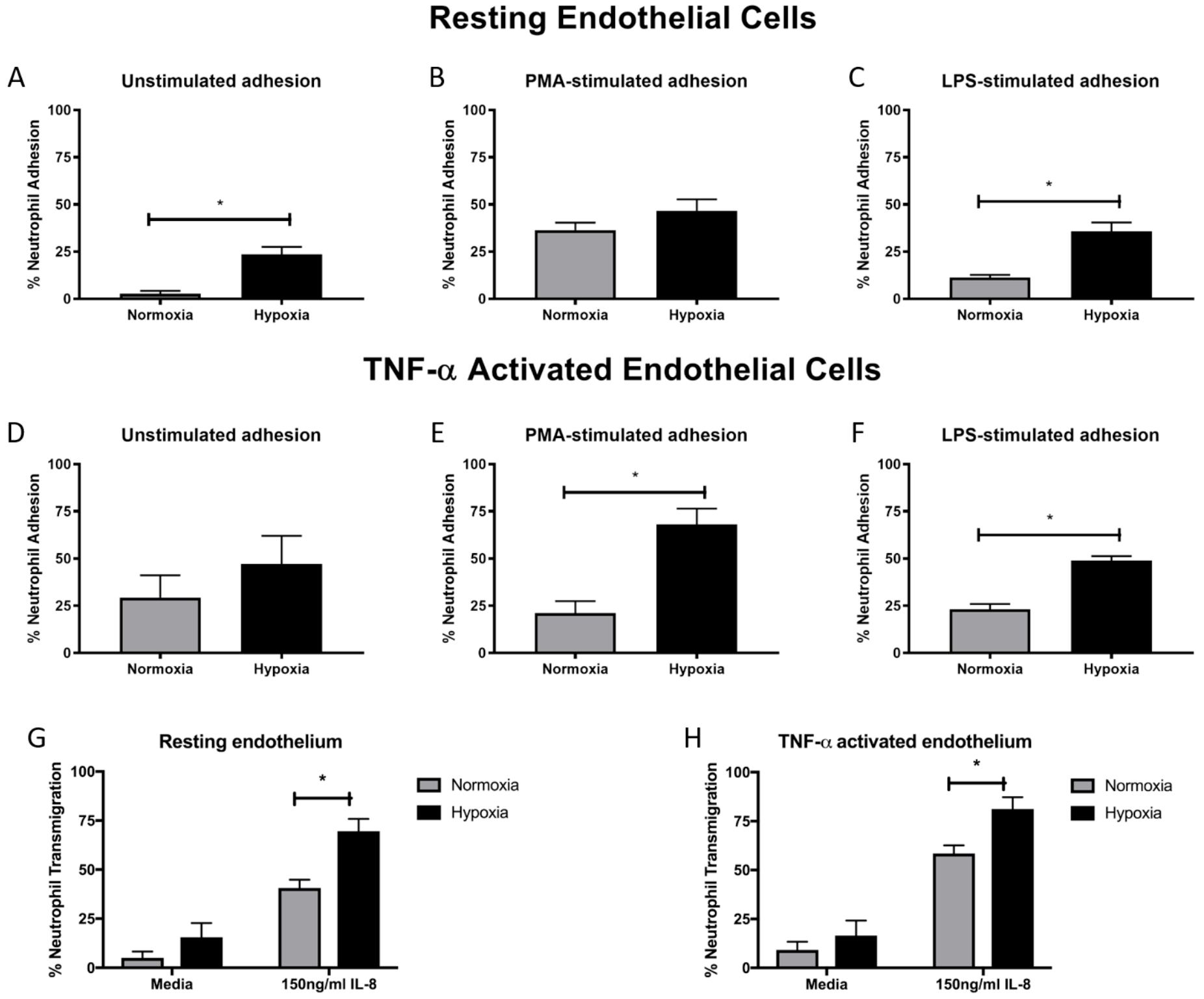
Hypoxia enhances neutrophil adhesion and trans-endothelial migration. BCECF-AM labelled neutrophil adhesion to endothelial monolayers over 30 minutes was assessed under normoxia (21% oxygen) and hypoxia (1% oxygen). We first examined neutrophil adhesion to resting HUVEC using **(A)** unstimulated neutrophils, **(B)** neutrophils stimulated with 20nM PMA and **(C)** cells stimulated with 100ng/ml LPS. Next, we examined the effects of hypoxia upon neutrophil adhesion to activated HUVEC, which had been stimulated with 10ng/ml TNF-α for 24 hours prior to experimentation. We evaluated **(D)** unstimulated neutrophil adhesion, **(E)** neutrophil adhesion in response to 20nM PMA and **(F)** neutrophil adhesion in response to 100ng/ml LPS. Finally, the effects of hypoxia upon trans-endothelial migration of CellTracker™ Green labelled neutrophils over 90 minutes was evaluated. **(G)** Neutrophil transmigration across resting endothelial monolayers was measured under both normoxia and hypoxia in the absence or presence of 150ng/ml IL-8. **(H)** HUVEC were stimulated with 10ng/ml TNF-α for 24 hours. Neutrophil trans-endothelial migration was subsequently measured in the absence or presence of 150ng/ml IL-8 under normoxic or hypoxic conditions. Data are presented as the mean and SEM from three independent experiments and analysed by a Wilcoxon matched pairs test or two-way ANOVA with a Dunnett’s multiple comparison test. *= p<0.05. Abbreviations: HIF, hypoxiainducible factor; PMA, phorbol 12-myristate 13-acetate.

Next, we evaluated neutrophil trans-endothelial migration in the absence and presence of IL-8, which has been shown to induce neutrophil migration. Basal transmigration (in the absence of IL-8) across both resting and TNF-α activated endothelium was unaffected by hypoxia. In contrast, in the presence of IL-8, hypoxia enhanced neutrophil trans-endothelial migration across both resting and TNF-α activated endothelium (p<0.05) (fig. 3G, H).

### Hypoxia increases expression of neutrophil β_2_ integrins, but not β_1_ integrins

Given the role of integrins in leukocyte extravasation and reports documenting reduced NETosis following integrin blockade^34^, we assessed surface integrin expression. Whilst α_L_ expression was unaffected (fig. 4A), significantly higher levels of α_M_ (3.1-fold increase, p<0.001) and α_X_ (1.6-fold increase, p<0.01) were observed under hypoxia (fig. 4B, C). There were no significant differences in β_2_ expression (fig. 4D). Hypoxia did not have an effect upon α_1_, α_4_, α_5_, or β_1_ integrin subunit expression (fig. 4E-H).

**Figure 4:**
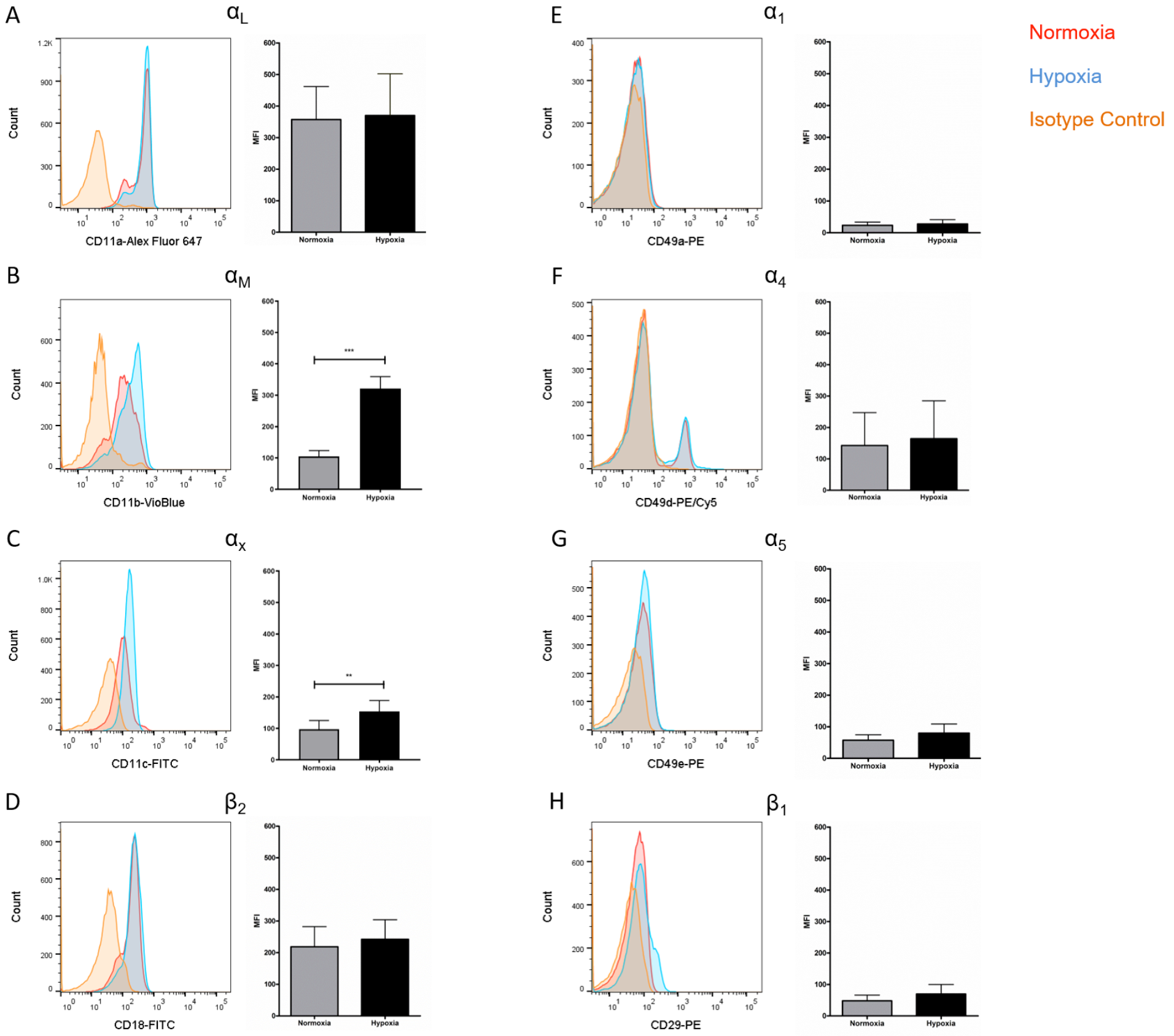
Hypoxia increased neutrophil α_M_ and α_X_ integrin subunit expression. Neutrophil integrin expression was examined following culture under normoxia (21% oxygen) or hypoxia (1% oxygen). Flow cytometry was used to assess expression of the integrin subunits: **(A)** α_L_, **(B)** α_M_, **(C)** α_X_, **(D)** β_2_, **(E)** α_1_, **(F)** α_4_, **(G)** α_5_ and **(H)** β_1_. Data are presented as the mean and SEM of neutrophils isolated from 7 different donors and analysed by Wilcoxon matched pairs test. **= p<0.01, ***= p<0.001. Abbreviations: HIF, hypoxia-inducible factor.

### NETosis is induced by α_M_β_2_ integrin activation

Given reports of reduced NETosis following integrin inhibition and our data showing increased α_M_β_2_ and, to a lesser extent, α_X_β_2_ integrin expression, we tested whether integrin-mediated adhesion activated NETosis in our model. As expected, no NETs were observed in unstimulated neutrophils, whilst PMA stimulation induced the externalisation of histone H3 (fig. 5A, B). Cation chelation through the use of EDTA, which abolishes integrin-mediated adhesion, suppressed PMA-induced NETosis (fig. 5C). Neutrophils stimulated with leukadherin-1 (LA-1), a compound that specifically activates the α_M_β_2_ integrin, produced histone H3 staining that was similar to PMA-stimulated cells (fig. 5D). Finally, global integrin activation by means of manganese chloride treatment, induced some histone H3 externalisation that could also be suppressed with EDTA treatment (fig. 5E, F). These results confirmed that α_M_β_2_ integrin activation can induce NETosis.

**Figure 5:**
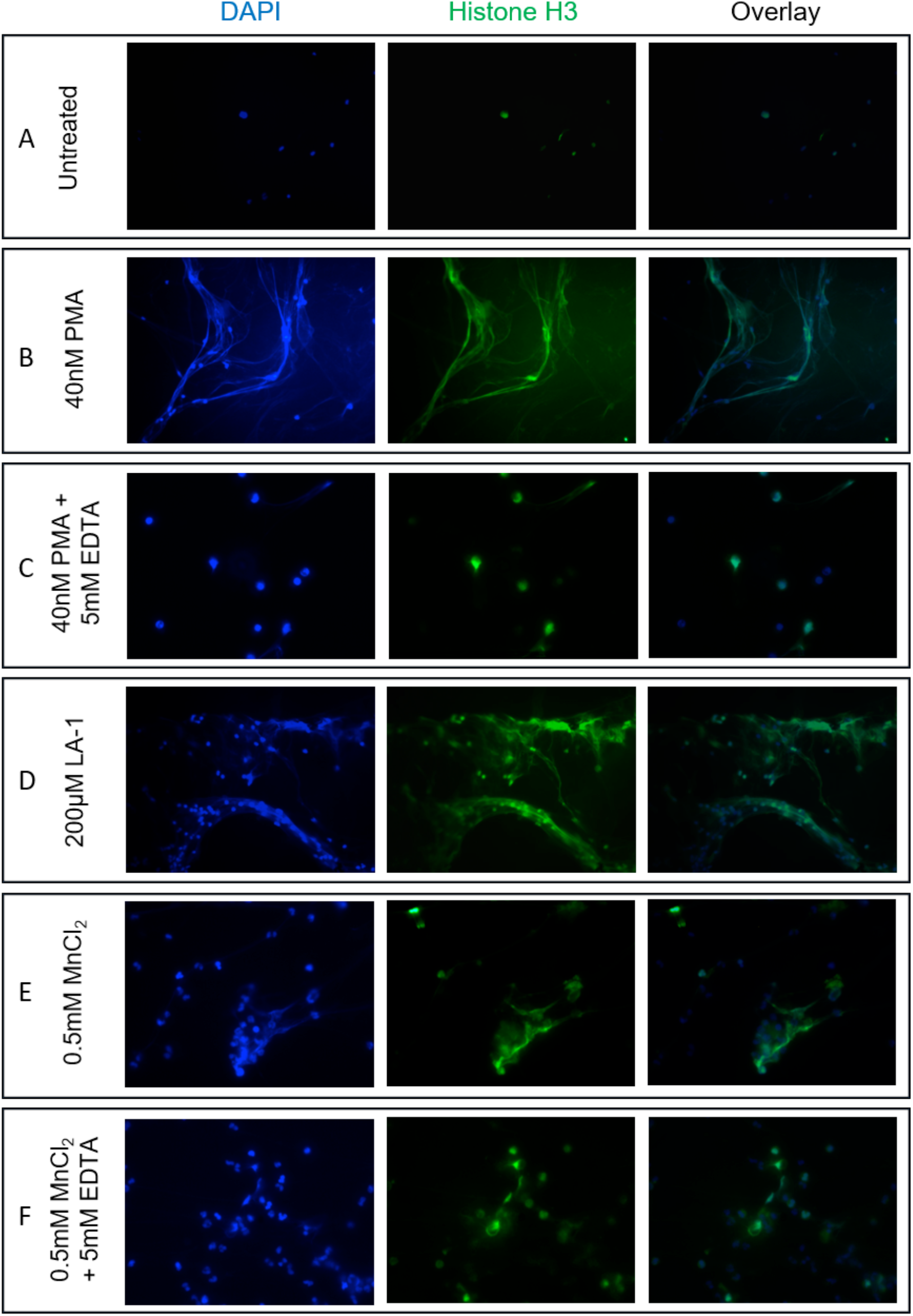
NETosis can be induce by neutrophil α_M_β_2_ integrin activation. The effects of cation-dependent integrin activation upon NETosis was examined visually by immunofluorescence. **(A)** Untreated neutrophils displayed punctate nuclear staining. **(B)** 40nM PMA stimulation induced DNA externalization. **(C)** PMA-induced NETosis could be mitigated by the addition of 5mM EDTA (cation chelator). We subsequently examined whether cationdependent integrin activation could induce NETosis. **(D)** Stimulation with 200μM leukadherin-1, a α_M_β_2_-specific integrin activator, also induced NETosis similar to PMA stimulation **(E)** Stimulation with 0.5mM MnCl_2_ induced some DNA externalisation. **(F)** DNA externalisation was suppressed following cation chelation with EDTA. Abbreviations: LA-1, leukadherin-1; Mn, manganese.

### BAL from patients with ILD have more NETs than control BAL

Having initially noted evidence of cell-specific hypoxia in ILD lung biopsies and demonstrating that hypoxia enhances neutrophil adhesion, transmigration and NETosis, we hypothesized that aberrant neutrophil activation may also be a feature of the ILD lung. To test this hypothesis, we generated slides with BAL and stained with DAPI to identify cellular DNA. Interestingly, whilst control BAL neutrophils displayed punctate DAPI staining (fig. 6A), we observed evidence of NETosis in ILD-BAL neutrophils (fig. 6B, C). We then obtained BAL from 11 ILD patients (ILD-BAL) and 7 non-ILD controls (control BAL) and quantified levels of cell-free DNA. We found ILD-BAL had 5.5-fold greater cell-free DNA content compared to control BAL (p<0.01) (fig. 6D). Cell-free DNA content positively correlated with neutrophil counts (% of total cells) isolated from ILD-BAL (p=0.0075) (fig. 6E), but not in control BAL (fig. 6F). To verify that these were NETs, we also examined BAL for the presence of MPO-citrullinated histone H3 complexes. Similar to total cell-free DNA, we observed significantly greater values in ILD-BAL, indicating greater levels of NETosis (fig. 6G).

**Figure 6:**
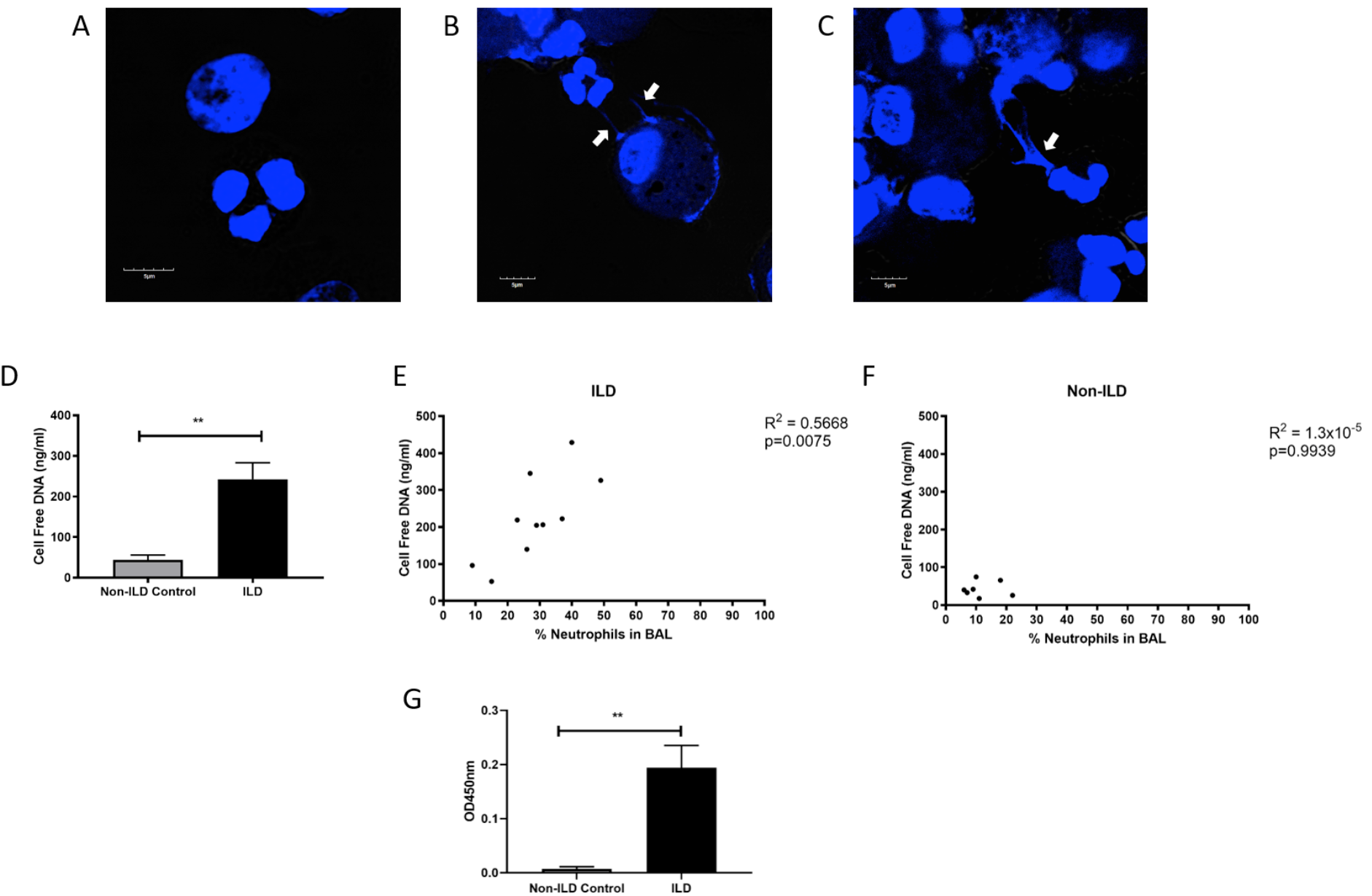
BAL isolated from ILD patients contain more NETs and NETosing neutrophils. BAL from ILD patients and non-ILD controls was examined for evidence of neutrophil activation. First, BAL fluid was assessed for the presence of NETs by **(A)** Quant-iT™PicoGreen^®^ dsDNA assay (Thermo Fisher, UK) and **(B)** an optimised capture ELISA detecting MPO-citrullianted histone H3 complexes. In both assays, BAL fluid from ILD patients has significantly more NETs compared to non-ILD controls. BAL cells were isolated and cytospin slide generated. Differential cell counts were made and the proportion of BAL neutrophils calculated. **(C)** Cell-free DNA content positively correlated with the percent neutrophils in the BAL of ILD patients, whilst **(D)** no correlation was seen in non-ILD controls. We finally stained cytospin slides with DAPI to visualize DNA. **(E)** Non-ILD control samples displayed punctate nuclear staining. **(F, G)** ILD samples, shown here from two patients, displayed evidence of DNA externalisation (indicated by white arrows). Data are presented as the mean and SEM from 11 ILD patients and 7 non-ILD controls and analysed by either a Mann-Whitney test or two-tailed Pearson correlation coefficients.**= p<0.01. Abbreviations: BAL, bronchoalveolar lavage; ILD, interstitial lung disease.

## Discussion

Neutrophil dysfunction and aberrant activation have been implicated in the pathology of numerous diseases including autoimmune rheumatic diseases^37–41^ and cancer^42–45^. More recently NETosis, essential for robust immune defense against pathogens, has also been linked to increased immunopathology in patients with autoimmune rheumatic disease^46–49^ and cancer^50–52^. The precise mechanism neutrophils contribute to ILD pathogenesis, however, is less characterised. Early work from the 1980s began to explore whether neutrophils might contribute to IPF pathology^11–14^, however this avenue of research lost momentum. Since then, reports have associated increased neutrophil migration and activation with severe pulmonary disease both in animal models^23,53^ and man^16,20,21^. We report, for the first time, that neutrophils and endothelial cells in ILD lung biopsies display HIF-1α expression and provide evidence for NETosis in the ILD lung. Given the profound effects hypoxia exerts upon neutrophil survival and function^27,28^, these findings led us to investigate whether hypoxia affects neutrophil extravasation and activation, thus contributing to ILD pathology.

Integrins are adhesive molecules that enable leukocytes to interact with their external environment. Similar to a previous report^54^, we found increased β_2_ integrin expression in neutrophils under hypoxia, but specifically found increased α_M_ and, to a lesser extent, α_X_ integrin subunit expression. Interestingly, the α_M_β_2_ and α_X_β_2_ integrins also function as complement receptors, which may be relevant to ILD pathology given that increased levels of complement C3a and C5a and roles for their receptors have been reported in IPF^55,56^. Moreover, studies using the bleomycin-induced mouse model of IPF highlight roles for both C3 and C5 in pulmonary fibrosis^57,58^. Upregulation of β_1_ integrins has been described under hypoxia^59^, however, there are no reports assessing expression in neutrophils. A lack of effect may be explained by the relatively low β_1_ integrin expression in human neutrophils. Taken together, the evidence indicates that neutrophils predominately engage via β_2_ integrins, a mechanism which is enhanced under hypoxia.

Early studies demonstrated that hypoxia enhances neutrophil adhesion to endothelial cells^60^, epithelial cells^61^, and trans-epithelial migration^62^. In support of these findings, our results show altered function of healthy neutrophils with increased neutrophil adhesion and trans-endothelial migration under hypoxia. In addition, we report that hypoxia enhances NETosis. Given that the gold standard markers or methods of detection for NETosis have not been established, we used several different techniques to confirm the release of NETs by cultured neutrophils: the co-localisation of nuclear DNA and histone H3 complexes by immunostaining, confocal imaging of DNA, measuring cell-free DNA, and the detection of neutrophil-derived proteins (MPO) and citrullinated histone H3 complexes. Our observations complement studies in the literature showing that pharmacological HIF-1α stabilisation enhances NETosis and inhibition of HIF-1α reduces NETs and bactericidal activity^29,30^. Taken together, our data, alongside published reports, indicate that hypoxia may drive the anti-bacterial effects of neutrophils through promoting NETosis.

We observed tissue-specific HIF-1α expression in ILD lung tissue, mainly restricted to pulmonary endothelial cells and neutrophils, with only minimal upregulation in areas of epithelium and fibrosis, which may hold pathological relevance. Previously, markers of hypoxia have been variably reported in the epithelium of patients with IPF. Several authors have found HIF-2α and CA-IX within the IPF fibrotic reticulum and HIF-1α in the overlying epithelium with IHC^7^, (albeit sometimes in a single patient^8^). HIF-1α is more readily found in the mouse bleomycin model of PF raising the question of differences between the two species and insults^63^.

Whilst epithelium-specific HIF-1α deletion has no effect upon radiation-induced enteritis, mice with endothelium-specific HIF-1α deletions present with reduced intestinal damage^64^. HIF-1α is known to contribute to the pathology of pulmonary hypertension^65,66^, with some work specifically interrogating endothelial HIF signalling^67^. Neutrophilic inflammation has also been associated with pulmonary hypertension^68^, and believed to drive angiogenesis via NETosis^69^. Therefore, the pulmonary pathology in ILD patients may in part be attributed to endothelial and neutrophil HIF-1α expression enhancing neutrophil recruitment and NETosis within the lung. In this paradigm, enhanced NETosis would initiate angiogenic signals and drive lung pathology. Interestingly, the model of neo-angiogenesis underlying ILD pathology has attracted interest, and the powerful angiogenic inhibitor, nintedanib, shown to have therapeutic benefits in a range of fibrotic ILDs^70^.

Our findings of hypoxia driving NETosis compliments the increasing evidence that NETs may play a role in many chronic lung diseases^71^, including ILD by stimulation of fibroblasts^17^. In PF, we propose that elevated NETosis may cause epithelial cell damage, dysfunction and death, drive innate and adaptive immune cells activation, and promote a pro-fibrotic environment that ultimately facilitates the progression of pulmonary fibrosis.

In conclusion, we report that the ILD lung contains molecular features of hypoxia, mainly localized to neutrophil and endothelial cells, which may contribute to disease pathology. Hypoxia enhanced neutrophil β_2_ integrin expression, which translated to augmented adhesion and migration across endothelial cells, and NETosis. Our findings are further supported by *ex vivo* demonstration of NETs within the human fibrotic lung. Taken together, our work begins to elucidate a potential role of hypoxia in driving neutrophil recruitment and activation within the airspace to promote a pro-fibrotic environment. These findings offer a rationale for future translational medicine exploration of a novel HIF-1α-Integrin-NETosis axis as a potential therapeutic target in fibrotic ILD.

## Methods

### Bronchoalveolar lavage

Fibre-optic bronchoscopy with BAL was performed in line with the American Thoracic Society guidelines^72^. BAL fluid was frozen for later NET analysis. None of the patients undergoing bronchoscopy had any infections at the time of procedure.

### Immunohistochemistry (IHC)

Lung biopsy specimens were collected as part of routine clinical care. Ethical approval was given by the UK National Research Ethics Committee (13/LO/0900). IHC was performed using the automated Bond-Max system (Leica Biosystems Ltd., Newcastle) with 4μm FFPE sections. HIF-1α (clone EP1215Y, Abcam, 1:600 dilution), MPO (polyclonal, Dako, 1:300 dilution) or NE (clone NP57, Dako, 1:100 dilution) was incubated in Epitope Retrieval Solution 2 for 20 minutes and stained using the 30,20,20 protocol. Test antibodies were controlled for using species- and isotype-matched control antibodies. Slides were scanned on a Nanozoomer Digital Slide Scanner and images analyzed using NDP viewer software (Hamamatsu Corportation). A “blinded” reviewer analysed five randomly selected areas from each subject. Representative images were chosen from those selected.

### Neutrophil isolation

Neutrophils were isolated as previously described^73^. In brief, neutrophils were isolated by Percoll density centrifugation from sodium citrate anticoagulated blood obtained by informed consent from healthy volunteers. Neutrophils were diluted to 2×10^6^ neutrophils/ml in phenol-free RPMI (Thermo Scientific, UK) supplemented with 10% FBS (Thermo Scientific, UK) and 2mM L-gluatamine (Lonza, UK).

### Endothelial cell culture

Human umbilical cord vein endothelial cells (HUVEC) (Lonza, Switzerland) were cultured in endothelial growth media 2 supplemented with 10% FBS (Thermo Scientific, UK) and 2mM L-glutamine (Lonza, UK) and used at passage 5. For endothelial activation, HUVEC were treated with 10ng/ml TNF-α (R&D Systems, UK) for 24 hours prior to experimentation.

### Hydrogen peroxide generation

H_2_O_2_ generation was measured as previously described^73^. Briefly, neutrophils were cultured under normoxia or hypoxia for 1 hour before addition of HRP (Sigma, UK) and Amplex^®^ UltraRed (Invitrogen, UK). H_2_O_2_ generation in response to PMA was recorded using a FLUOstar Omega microplate reader (BMG Labetech, Germany) and rates (expressed in nM/sec) determined using Omega Mars Analysis software (BMG Labtech, Germany).

### NETosis quantification

NETs was quantified using the Quanti-iT™ PicoGreen^®^ dsDNA kit (Invitrogen, UK). Supernatants were also tested using a capture ELISA. Streptavidin-coated plates (Fisher Scientific, UK) were coated with an anti-MPO capture antibody (Abcam, UK) overnight at 4°C and blocked with 0.5% bovine serum albumin for 1 hour at 37°C. Neutrophil supernatants were incubated for 2 hours at 37°C. Further 1 hour incubations were performed with an anti-citrullinated histone H3 detection antibody (Abcam, UK) and HRP-conjugated secondary antibody (Dako, UK). SureBlue TMB Microwell Peroxidase Substrate (KPL, UK) was then added and incubated in the dark at 37°C for 20 minutes and then stopped by the addition of TMB stop solution (KPL, UK). Absorbance was read at 450nm using a Tecan GENios Spectra FLUOR plate reader (Tecan UK Ltd., UK).

### NET immunofluorescence

NETs were stained for immunofluorescence microscopy as described^73^. In brief, 5×10^5^ neutrophils were added to coverslips, stimulated, and then fixed with 4% PFA. Coverslips were blocked and sequentially incubated with an anti-histone H3 antibody (Abcam, UK) and Alexa Fluor^®^ 488-conjugated goat anti-rabbit IgG secondary antibody (Life Technologies, UK). Coverslips were washed, mounted, and sealed using with ProLong™ Gold antifade mountant with DAPI (Invitrogen, UK). Slides were visualized using a Zeiss Axio Imager.A1 inverted fluorescence microscope (Zeiss, Germany) and images analysed using Image J.

### BAL confocal immunofluorescence

BAL fluid was filtered using a 40μm cell sieve. BAL cells were pelleted, counted and 1×10^5^ viable cells were used to produce cytospin slides (Thermo Shandon Cytospin 3, Thermo Scientific). Cytospin slides were fixed in 4% PFA, washed, and blocked overnight in blocking solution (10%goat serum/1% BSA/2mM EDTA/HBSS/0.1% Tween-2). Slides were then washed and incubated with DAPI (Sigma, UK) diluted in blocking buffer for 1 hour. Stained slides were then washed, mounted, sealed and visualised using an Olympus inverted fluorescence confocal microscope and analysed using Image J.

### Neutrophil integrin expression

Cell surface expression of neutrophil integrins (α_1_ β_1_, α_4_β_1_, α_5_β_1_, α_L_β_2_, α_M_β_2_ and α_X_β_2_) was evaluated by flow cytometry. Following isolation and culture under either normoxia or hypoxia, neutrophils were washed and resuspended in a sodium HEPES buffer (20mM HEPES, 140mM NaCl, 2mg/ml glucose, 0.3% BSA). Cells were then stained using integrin subunit specific antibodies or appropriate isotype control for 30 minutes at room temperature. Stained cells were then washed twice, fixed and assessed using a FACS Verse (BD Biosciences, UK). Data was analysed using FlowJo (TreeStar Inc., UK).

### Neutrophil adhesion

HUVEC were cultured in 96-well black tissue culture plates (Thermo Scientific, UK). 24 hours prior to experimentation, HUVEC were subjected to normoxia or hypoxia in the absence or presence of 10ng/ml TNF-α. Neutrophil adhesion in response to 20nM PMA or 100ng/ml LPS were measured as previously described^73^. Briefly, neutrophils were cultured under normoxia or hypoxia for 1 hour, then labelled with 2’,7’-bis-(2-carboxyethyl)-5-(and-6)-carboxyfluoresceinacetoxymethyl ester (Life Technologies, UK). Neutrophils were then added to wells under normoxia or hypoxia. Fluorescence was measured using a Tecan GENios Spectra FLUOR plate reader (Tecan UK Ltd., UK). Adhesion was calculated by comparing the fluorescence of washed wells to initial fluorescence.

### Neutrophil trans-endothelial migration

Trans-endothelial migration assays were performed as previously described^73^. In brief, HUVEC were grown on transwell inserts (Millipore, UK). 24 hours prior to experimentation, HUVEC were cultured under normoxia or hypoxia in the absence or presence of 10ng/ml TNF-α. Neutrophils were cultured under normoxia or hypoxia for 1 hour and then labelled with CellTracker (Invitrogen, UK). 1×10^6^ neutrophils were added to the upper chamber of transwells and allowed to migrate in the absence or presence of 150ng/ml IL-8 in the lower chamber for 90 minutes. Percent transmigration was calculated by comparing the number of cells in the lower chamber and the number of neutrophils added to the upper chamber.

### Western blotting

Cell lysates (10μg protein) were resolved by electrophoresis and transferred to a polyvinylidene fluoride membrane (GE Healthcare, UK). Membranes were blocked for 1 hour in 5% skimmed milk/TBS/0.1% Tween-20 and incubated with primary antibodies (1:1000 dilution) overnight at 4°C. Membranes were then washed, incubated with HRP-conjugated secondary antibodies, and visualised using the Luminata Western HRP substrate system (Millipore, Ireland).

### Statistical analysis

Data were evaluated using GraphPad Prism. Data were tested for normality using a Kolmogorov-Smirnov test. In experimental data sets only comparing two groups, a Mann-Whitney test was performed or a Wilcoxon matched pairs test. In data sets with two variables, data were assessed by two-way ANOVA with a Dunnet’s multiple comparison test. Correlations were determined by two-tailed Pearson correlation coefficients. A p value below 0.05 was considered significant.

## Acknowledgements

We would like to acknowledge Professor Margaret Ashcroft and Dr Luke Thomas (University of Cambridge) for advice and input on HIF, Dr Andrew Smith and Professor Tony Segal (University College London) for their help with the neutrophil H_2_O_2_ generation assay, Professor Nancy Hogg (The Francis Crick Institute) for reagents and helpful discussions, and Dr Guiseppe Ercoli (University College London) who helped with confocal imaging.

