## supplementary table 1 for "Identification of a novel HIF-1α-α_M_β_2_ Integrin-NETosis axis in fibrotic interstitial lung disease"

**Supplementary Data**

| Sample | Differential Cell Count (%) | | | |
|---|---|---|---|---|
| | **Macrophage** | **Neutrophil** | **Lymphocyte** | **Eosinophil** |
| ILD01 | 59 | 29 | 5 | 6 |
| ILD02 | 47 | 27 | 8 | 19 |
| ILD03 | 47 | 40 | 3 | 10 |
| ILD04 | 35 | 49 | 10 | 7 |
| ILD05 | 62 | 23 | 2 | 14 |
| ILD06 | 79 | 9 | 12 | 1 |
| ILD07 | 59 | 31 | 8 | 2 |
| ILD08 | 66 | 26 | 4 | 5 |
| ILD09 | 69 | 26 | 4 | 1 |
| ILD10 | 60 | 37 | 2 | 0 |
| ILD11 | 68 | 22 | 1 | 8 |
| Non-ILD01 | 72 | 22 | 3 | 3 |
| Non-ILD02 | 81 | 11 | 8 | 0 |
| Non-ILD03 | 91 | 6 | 2 | 2 |
| Non-ILD04 | 92 | 6 | 3 | 0 |
| Non-ILD05 | 78 | 18 | 0 | 3 |
| Non-ILD06 | 86 | 10 | 3 | 1 |
| Non-ILD07 | 97 | 3 | 0 | 0 |

**Supplementary Table 1: Cellular compositions of bronchoalveolar lavage (BAL) fluid.** BAL fluid was obtained from patients undergoing diagnostic bronchoscopy. BAL cells were isolated by centrifugation, differentially stained with a Rapid Romanowsky Stain Kit (TCS Biosciences Ltd, UK) and counted using a conventional brightfield microscope.
